# Photoperiodic Effects on Monoamine Signaling & Gene Expression Throughout Development in the Serotonin & Dopamine Systems

**DOI:** 10.1101/2020.06.25.171470

**Authors:** Justin K. Siemann, Piper Williams, Turnee N. Malik, Chad Jackson, Noah H. Green, Ronald Emeson, Pat Levitt, Douglas G. McMahon

**Affiliations:** Vanderbilt University, Nashville, TN, 37232; Children’s Hospital of Los Angeles, Los Angeles, CA, 90027

**Keywords:** photoperiod, development, serotonin, dopamine, dorsal raphe nucleus, sensitive periods, monoamine content, gene expression, quantitative RTPCR, RNAScope

## Abstract

Photoperiod or the duration of daylight has been implicated as a risk factor in the development of mood disorders. The dopamine and serotonin systems are impacted by photoperiod and are consistently associated with affective disorders. Hence, we evaluated, at multiple stages of postnatal development, the expression of key dopaminergic (*TH*) and serotonergic (*Tph2, SERT,* and *Pet-1*) genes, and midbrain monoamine content in mice raised under control Equinox (LD 12:12), Short winter-like (LD 8:16), or Long summer-like (LD 16:8) photoperiods. Focusing in early adulthood, we evaluated the midbrain levels of these serotonergic genes, and also assayed these gene levels in the dorsal raphe nucleus (DRN) with RNAScope. Mice that developed under Short photoperiods demonstrated elevated midbrain *TH* expression levels, specifically during perinatal development compared to mice raised under Long photoperiods, and significantly decreased serotonin and dopamine content throughout the course of development. In adulthood, Long photoperiod mice demonstrated decreased midbrain *Tph2* and *SERT* expression levels and reduced *Tph2* levels in the DRN compared Short photoperiod mice. Thus, evaluating gene x environment interactions in the dopaminergic and serotonergic systems during multiple stages of development may lead to novel insights into the underlying mechanisms in the development of affective disorders.

## Introduction

Globally it is estimated that over 300 million individuals suffer from depression and 17 million Americans have reported experiencing at least one depressive episode^1,2^. Throughout adulthood the prevalence of depression is approximately 7%^2^. However, it is highest during early adulthood between the ages of 18-25 (13%)^2^ and in adolescence, between the ages of 12-17, the prevalence can reach similar rates peaking at 13%^3^. Interestingly, the prevalence of childhood depression is only approximately 2%, yet there is evidence that this and the overall incidence of mood disorders is beginning to steadily increase over the last 10 years^4^. Thus, childhood, adolescence, and early adulthood may represent vulnerable periods in which individuals may be more susceptible to the risk of developing depression^5–7^.

Exposures to environmental factors during sensitive periods of development have been associated with elevated risk for neurodevelopmental disorders later in adulthood^8^. To this point, studies have demonstrated significant development x environment effects resulting in increased prevalence for psychiatric disorders^9^. An important environmental factor, the duration of daylight or photoperiod, has been implicated in affective disorders in adolescence and adulthood^10–12^, and has been shown to have profound and lasting effects on mood-related behaviors^12–17^ Photoperiod has been linked to increased risks for psychiatric disorders^11,18–21^, can modulate monoamine turnover of serotonin and dopamine^22,23^, and associations between photoperiod and genes relevant to the serotonin and dopamine systems have revealed significant gene x environment effects observed during the winter and fall months, when the duration of daylight is lowest during the year^24^. Therefore, photoperiod appears to be a consistent environmental factor that may play a critical role in the development of affective disorders.

Rodent studies in the dopaminergic and serotonergic systems have shown that photoperiodic exposure significantly affects neuronal firing rate^15,32^, monoamine signaling^33–35^, and mood-related behaviors^17,36,37^, which can be sex-dependent^34,38^. Specifically, it has been shown that *developmental* photoperiod can program various aspects of the serotonin system^15,34^ Mice that developed under Long summer-like photoperiods demonstrate increased dorsal raphe (DRN) serotonin (5-HT) neuronal firing rate, elevated midbrain serotonin content, and reduced anxiety and depressive-like behaviors later in life in a melatonin receptor 1 (MT1R) dependent manner^15^. Our lab has recently shown that *prenatal* photoperiodic exposure results in enduring changes to DRN 5-HT neuronal activity, and there are critical temporal windows within *perinatal* development that are impacted by photoperiod resulting in enduring changes to monoamine signaling and affective behaviors during adolescence and early adulthood^34^.

The midbrain contains both dopamine and serotonin rich nuclei, is a critical brain region responsible for mood regulation, and disruptions in these circuits have been linked to depression^8,39,40^. Midbrain dopaminergic neurons have specifically been shown to regulate aspects of depressive-like behavior and the activity of these neurons may be important for resilience against depression^41,42^. Tyrosine hydroxylase (*TH*) is a key gene in the dopamine system as it is the rate-limiting enzyme for dopamine (DA) synthesis, alterations in this gene have been observed in patients with depression, and have led to investigations focused on targeting *TH* as a novel therapeutic treatment for mood disorders^43–45^. Importantly, it has been shown that photoperiod can modulate human gene expression levels of *TH* and the dopamine transporter (*DAT*) in the midbrain^29^.

The main center for serotonin synthesis and neuronal development is a structure within the midbrain, known as the dorsal raphe nucleus (DRN)^46^. Studies have demonstrated developmental x environment changes specifically in the serotonergic system as it pertains to mood related disorders^47–51^. Prior developmental work has shown that manipulations to serotonin (5-HT) receptors, the serotonin transporter, and environmental factors such as stress can dramatically alter serotonin synthesis, circuit formation along with anxiety and depressive-like behaviors that can persist throughout adulthood^52–61^. Tryptophan hydroxylase 2 (*Tph2*) is the rate limiting enzyme for serotonin synthesis, the serotonin transporter (*SERT*) is responsible for reuptake of serotonin back into presynaptic vesicles from the synaptic cleft, and the ETS transcription factor *Pet-1* is the main regulator for serotonin neuronal development and differentiation^62^. These three key 5-HT genes are highly expressed in serotonergic neurons and thus largely expressed in the midbrain and specifically the DRN^62^. Critically, developmental roles for these genes have been shown such that modulation of *Tph2, SERT* and/or *Pet-1* expression during prenatal or perinatal development can also result in vast molecular, circuit level, and behavioral changes relevant to mood disorders^63–67^.

In this study we evaluated the effects of photoperiod on midbrain dopamine (*TH*) and serotonin (*Tph2*, *SERT* and *Pet-1*) gene expression, midbrain monoamine content, and expression levels of these 5-HT genes specifically in the DRN. These assays were performed in the mouse during multiple sensitive periods of postnatal development, representing childhood, adolescence, and early adulthood^68^, which have been implicated in the development of mood disorders^7^, in order to investigate the role of photoperiod on the dopamine and serotonin systems during the course of development.

## Materials and Methods

### Animals and Housing

Male and female C3Hf^+/+^ mice were used as these animals are melatonin-producing and lack the retinal degeneration alleles of the parent C3H strain^69^. For developmental midbrain quantitative RTPCR and monoamine content experiments mice were raised under either an Equinox (Eq) (12 hours of light 12 hours of darkness), Long (L) (16 hours of light and 8 hours of darkness), or Short (S) (8 hours of light and 16 hours of darkness) photoperiod (**Figure 1 A-C).** We have used these photoperiods commonly in our previous work^15,34,70^ and these light-dark cycles (LD 16:8 and LD 8:16) mimic real-world photoperiods of summer-like (Long) and winter-like (Short) photoperiods at the high middle latitudes (e.g. equivalent to London, Paris, Berlin) and are experienced by significant portions of the human population. For all studies, mice were maintained continuously under these photoperiods from embryonic day 0 (E0) until they were assessed experimentally (**Figure 1**). Developmental midbrain quantitative RTPCR, measuring key dopamine (*TH*) and serotonin (i.e. *Tph2, SERT,* and *Pet-1*) genes, along with midbrain monoamine signaling assays were performed at P8, P18 (ranging from P17-P19), and P35 (ranging from P34-P37), representing perinatal development, early childhood, and adolescence in the mouse^68^ (**Figure 1**). For early adulthood midbrain RTPCR 5-HT gene experiments, mice developed and were raised from E0 to maturity under either an Equinox, Long, or Short photoperiod (**Figure 4 A-C)** and RNAScope measurements were evaluated from the DRN in mice that developed under either a Long or Short photoperiod (**Figure 4 B-C)**. Early adulthood quantitative RTPCR and RNAScope assays for serotonin related genes (i.e. *Tph2, SERT,* and *Pet-1*) were performed at postnatal day P50 (ranging from P50-P90), representing early adulthood in the mouse^71^ (**Figure 4**). All tissues were isolated at 1100-1300, the mid-day point on all light cycles. Mice were group housed and allowed access to food and water *ad libitum.* Ethical approval was obtained from the Vanderbilt University Institutional Animal Care and Use Committee and all experiments were performed in accordance with the Vanderbilt University Institutional Animal Care and Use Committee and National Institutes of Health guidelines.

**Figure 1.**
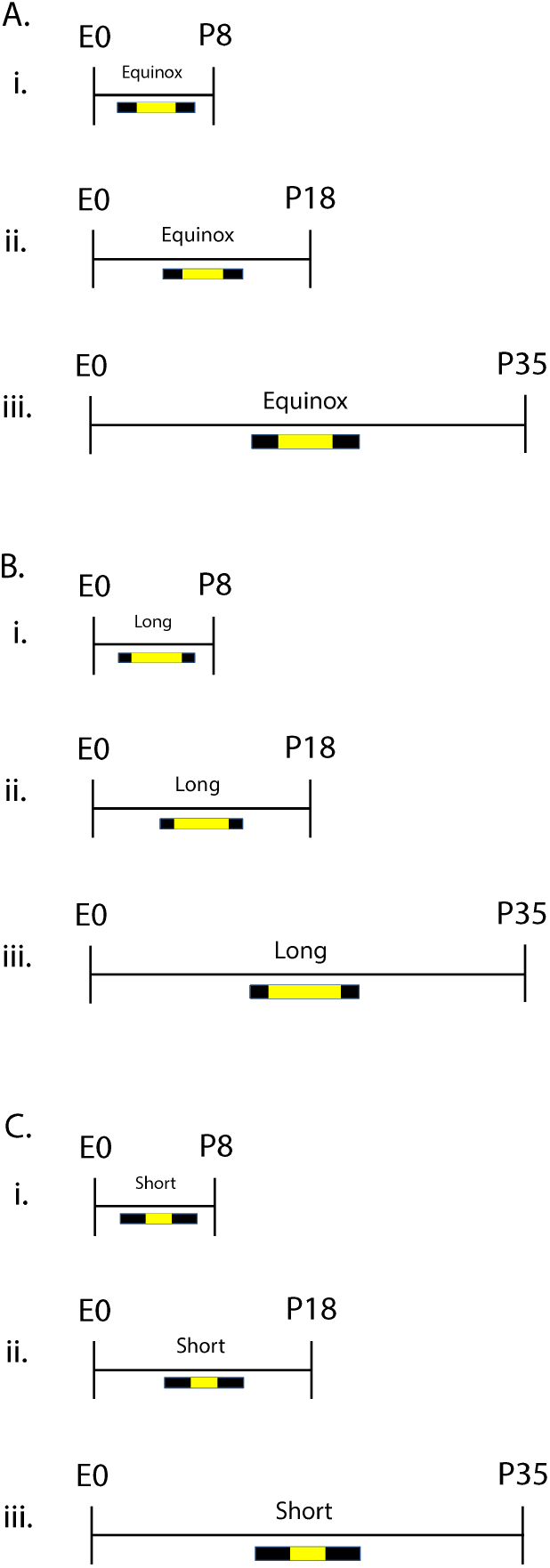
Photoperiod paradigm across development. For <u>developmental quantitative RTPCR and midbrain monoamine content experiments</u>, animals developed under either an **A)** Equinox LD 12:12, **B)** Long LD 16:8, or **C)** Short LD 8:16 photoperiod. Mice developed under these photoperiods from embryonic day 0 (E0) and were evaluated at postnatal days i) 8, ii) 18, or iii) 35 (i.e. P8, P18, P35). L = light, D = dark (i.e. LD 16:8 means Light for 16 hours, Dark for 8 hours).

### Developmental Midbrain Quantitative RTPCR Analysis

Mouse midbrains (n = 5-6 per group per age) were collected and RNA was isolated using TRIzol Reagent (Invitrogen) according to the manufacturer’s instructions. All RNA samples were treated with TURBO DNase (Invitrogen) to remove contaminating genomic DNA following manufacturer’s instructions. Reverse transcription was performed from isolated RNA using the QuantiTect Reverse Transcription Kit (Qiagen) containing a mixture of oligo-dT and random primers. Quantitative PCR (qPCR) reactions were set-up to include 1 μL of cDNA, 7 μL nuclease-free H20, 10 μL of TaqMan Universal PCR Master Mix (Applied Biosystems), 1 μL mouse β-actin primer-limited TaqMan Endogenous Control probe (ACTB VIC/MGB), and 1 μL target probe TaqMan probe (Mm00447557_m1 FAM-MGB). All samples were amplified for 36 cycles and relative *TH, Tph2, SERT,* and *Pet-1* expression were calculated using the ΔΔCt method^72^. Each sample was assayed in triplicate.

### Developmental Midbrain Monoamine Metabolism

Mouse midbrains (n = 11-14 per group and age) were dissected with clean razor blades from the inferior colliculus to the apex; −6.1mm to −4.1mm from Bregma. The tissue was placed in 1.5mL tubes and then frozen in liquid nitrogen and biogenic amine analysis was performed in the Vanderbilt Neurochemistry Core Laboratory. Briefly, samples were stored at −80°C, then the tissue was homogenized with a tissue dismembrator, in 100-750 μl of 0.1M TCA. This contained 10-2 M sodium acetate, 10-4 M EDTA, 5ng/mL isoproterenol (as internal standard) and 10.5 % methanol (pH 3.8). Samples were then spun in a microcentrifuge at 10000xg for 20 minutes with the supernatant being removed and stored at −80°C and the pellet were saved for protein analysis. Supernatant was then thawed and spun for 20 minutes and the samples of the supernatant were analyzed for biogenic amines. These amines were determined by a specific HPLC assay utilizing an Antec Decade II (oxidation: 0.4) (3mm GC WE, HYREF) electrochemical detector operated at 33°C. Twenty μl samples of the supernatant were injected using a Water 2707 autosampler onto a Phenomenex Kintex (2.6u, 100A) C18 HPLC column (100 x 4.60 mm), biogenic amines were eluted with a mobile phase consisting of 89.5% 0.1M TCA, 10-2 M sodium acetate, 10-4 M EDTA and 10.5% methanol (pH 3.8) and the solvent was then delivered at 0.6 mL/min using a Waters 515 HPLC pump. Utilizing this HPLC solvent the following biogenic amines eluted were: Noradrenaline, Adrenaline, DOPAC, Dopamine, 5-HIAA, HVA, 5-HT, and 3-MT and HPLC control and data acquisition were managed by Empower software. Methods were the same as those previously published^34^.

### Early adulthood quantitative RTPCR 5-HT Gene expression analysis

Mouse midbrains (n = 6 per group) were removed, frozen in a 1.5 mL tube in liquid nitrogen, and samples were stored at −80°C until RNA extraction. Total RNA was extracted using a Qiagen RNeasy mini kit (Qiagen Inc., Valencia, CA, USA, Cat. No. 74104), measured by a Nanodrop system (ThermoScientific), and reverse-transcribed (~200ng) into cDNA using the QuantiTect Reverse Transcription Kit (Qiagen Inc., Valencia, CA, USA, Cat. No. 205311). qRT-PCR reactions were performed in 20 μL total volume with 2 μL cDNA, 10μL of SsoAdvanced SYBR Green Supermix (Bio-Rad, Hercules, CA), 6μL sterile water and 1μL of 300 nM intron-spanning gene specific forward and reverse primers in a Bio-Rad CFX96 Real-Time System (Bio-Rad, Hercules, CA, USA). Quantification of transcript levels were performed by comparing the threshold cycle for amplification of the unknown to those of six concentrations of standard cDNAs for each respective transcript, then normalizing the standard-calculated amount to hypoxanthine guanine phosphoribosyl transferase (*Hprt*) in each sample. Each sample was assayed in duplicate.

### Early Adulthood RNAScope and Confocal Microscope Imaging

Mouse (n = 6 per group) whole brain tissue was collected and submerged into isopentane for 25 seconds, placed in crushed dry ice for approximately 1 minute, and stored in a 50mL falcon tube at −80°C. 16 μm sections were prepared per animal targeting the dorsal raphe (DRN) from −5.5mm to −5.75mm from Bregma. Tissue sections were then stained with RNAScope in situ hybridization, targeting the transcripts *Tph2, SERT,* and *Pet-1* with fluorescent dyes Alexa488, Atto550, and Atto647, respectively. For each animal, the section that included the area of interest (i.e. dorsal raphe nucleus) was determined based on serotonergic cell quantity and distribution. 512 x 512 pixel 2-dimensional scans of the three independent channels were collected using a Zeiss LSM510 confocal scanning microscope (Carl Zeiss Microscopy Gmbh, Jena, Germany) at 20X magnification. Due to the heterogeneity in the dorsal raphe nucleus (DRN), we focused on the ventromedial portion of the DRN. 75-95% of the cells in this portion of the DRN across the rostro-caudal axis are serotonergic and express the 5-HT1A receptor^73^. Integrated density for cell fluorescence and total ROI (i.e. DRN) fluorescence levels, mean cell fluorescence levels, mean total ROI (i.e. DRN) fluorescence, and total number of cells expressing each transcript were determined using the software program ImageJ (NIH, Bethesda, Maryland). Integrated density can be defined as the product of mean fluorescence and area. The draw tool in ImageJ was used in order to define each individual cell and ROI (i.e. the dorsal raphe nucleus).

### Data Analysis

Prism 8 (Graphpad Software Inc., La Jolla, CA) was used for all statistical analyses. Statistical significance was determined by either two-way ANOVA (for developmental midbrain RTPCR and monoamine signaling assays), one-way ANOVA (for early adulthood RTPCR 5-HT gene experiments), or paired t tests (for early adulthood DRN RNAScope experiments) with a p value less than 0.05 considered significant. Paired twotailed t tests were used for early adulthood RNAScope experiments because samples from each photoperiod were sectioned and stained together (i.e. one Long photoperiod animal and one Short photoperiod animal were sectioned and stained together on the same day). All post hoc analysis with Holm-Sidak’s multiple comparison tests were performed and standard error of the mean was used for all experiments unless otherwise specified.

## Results

### Perinatal *TH* Expression is Increased Under Short Winter-like Photoperiodic Conditions in the Midbrain

We investigated the role of developmental photoperiod on the expression of key dopamine (*TH*) and serotonin (i.e. *Tph2, SERT,* and *Pet-1*) genes in the midbrain through multiple stages of development. By utilizing quantitative RTPCR we evaluated the expression levels of these dopamine and serotonin genes at postnatal days (P) 8, 18, and 35 in mice that continuously developed under control Equinox, Long summer-like, or Short winter-like photoperiod conditions from E0 **(Figure 1)**. When measuring *TH* levels and utilizing a two-way ANOVA we found a significant main effect of photoperiod (p < 0.0001; F (2, 45) = 12.98), a main effect of age (p = 0.0037; F (2, 45) = 6.350), and an interaction effect (p = 0.0152; F (2, 45) = 3.455) **(Figure 2)**. At P8, significant Holm-Sidak’s multiple comparison post hoc tests were revealed between Short and Equinox (p < 0.0001), and between Short and Long photoperiods (p < 0.0001). No significant multiple comparison effects were observed between photoperiods at either P18 or P35 for midbrain *TH* expression levels. Trend level differences at P18 were observed between Short and Equinox (p = 0.0793), however these comparisons did not reach statistical significance. With quantitative RTPCR, we found that *TH* expression levels in the midbrain are significantly elevated during perinatal development (P8) and decrease throughout the course of development, normalizing by early adolescence (P35).

**Figure 2.**
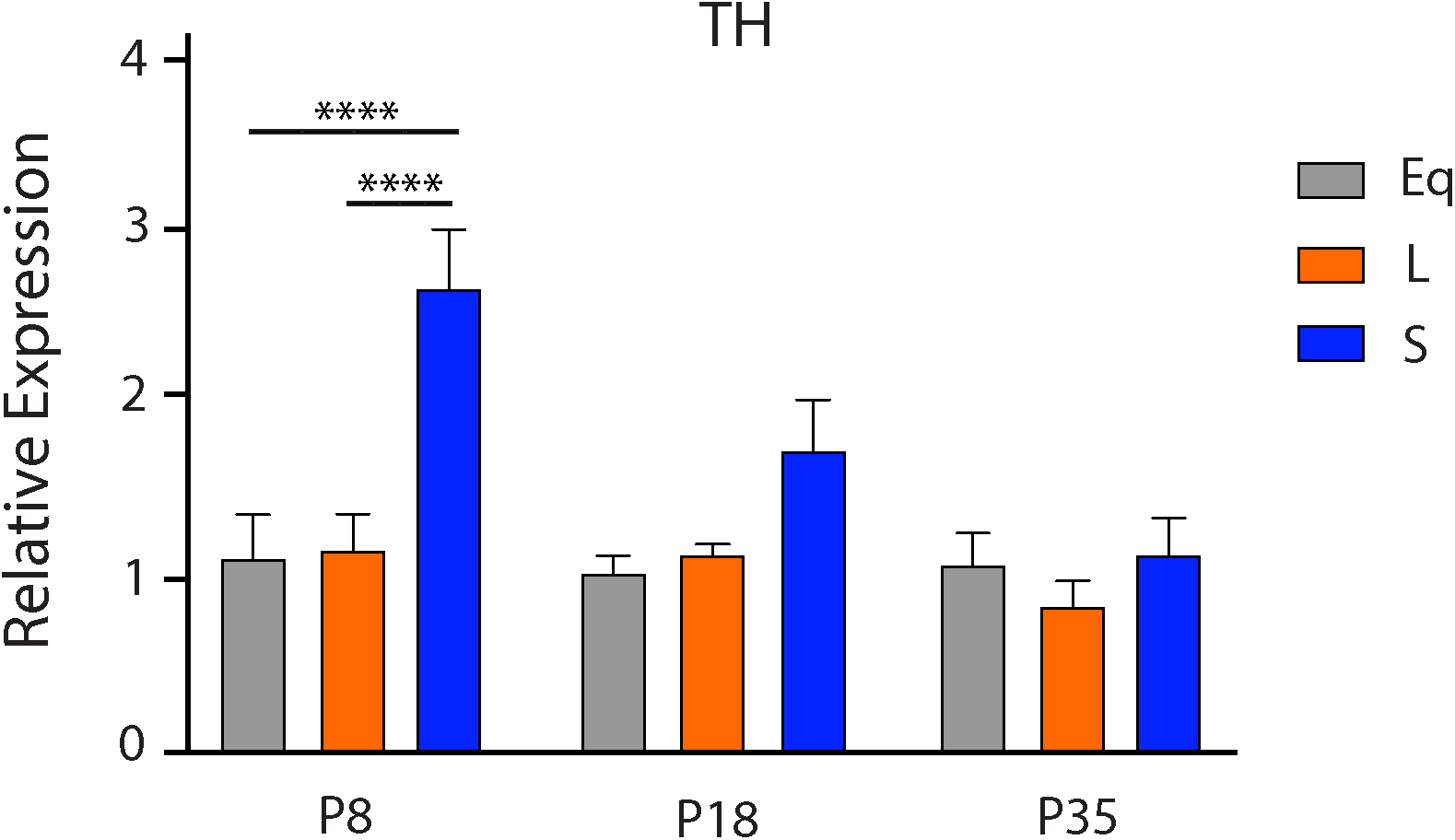
Perinatal Short photoperiod significantly increased midbrain *TH* levels. Mice developed under either Equinox (Eq), Long (L), or Short (S) photoperiods and *TH* expression levels were measured at three time points: P8, P18, and P35. The significant levels were as follows: (**** = p < 0.0001). Reference gene for relative expression was β-actin.

We also evaluated the effects of photoperiod throughout development on expression levels of key serotonergic genes (i.e. *Tph2, SERT,* and *Pet-1).* We found no significant main effects of photoperiod for these genes throughout development. When measuring *Tph2* levels and utilizing a two-way ANOVA we found a non-significant main effect of photoperiod (p = 0.6330; F (2, 41) = 0.4625) and a significant main effect of age (p = 0.0281; F (2, 41) = 3.901) **(Supplementary Figure 1)**. When measuring *SERT* levels and utilizing a two-way ANOVA we found non-significant main effects of photoperiod (p = 0.1913; F (2, 45) = 1.716) and of age (p = 0.0717; F (2, 45) = 2.796) **(Supplementary Figure 2)**. At P35, significant Holm-Sidak’s multiple comparison post hoc tests were revealed between Equinox and Long (p = 0.0179), and between Equinox and Short photoperiods (p = 0.0078). *Pet-1* gene expression levels were so low in the midbrain that they were undetectable. Overall, photoperiod did not impact expression levels in serotonin-related genes in the midbrain throughout development.

### Short Winter-like Photoperiod Exposure Results in Decreased Midbrain Serotonin & Dopamine Content at Multiple Development Periods

We were also interested in determining the effects of photoperiod on midbrain monoamine content during sensitive periods of postnatal development. We evaluated mice that were raised and developed under either control Equinox, Long summer-like or Short winter-like photoperiods from E0 and measured monoamine concentrations at P8, P18, and P35 **(Figure 1)**. We evaluated serotonin (5-HT) and dopamine (DA) content along with their main metabolites, 5-hydroxyindoleacetic acid (5-HIAA) and 3,4-Dihydroxyphenylacetic acid (DOPAC), respectively. For 5-HT content, using a two-way ANOVA, we found a significant main effect of photoperiod (p = 0.0166; F (2, 102) = 4.269), a main effect of age (p < 0.0001; F (2, 102) = 412.8), and an interaction effect (p < 0.0001; F (4, 102) = 8.897) **(Figure 3A)**. At P18, a significant Holm-Sidak’s multiple comparison post hoc test was observed between Long and Equinox (p = 0.0035) mice, and at P35, significant Holm-Sidak’s multiple comparison post hoc tests were observed between Long and Short (p = 0.0001), and Equinox and Short (p = 0.0014) photoperiod groups. For 5-HIAA concentrations, utilizing a two-way ANOVA, a significant main effect of photoperiod (p = 0.0183; F (2, 102) = 4.164), a main effect of age (p < 0.0001; F (2, 102) = 163.1), and an interaction effect (p = 0.0116; F (4, 102) = 3.411) were found **(Figure 3B)**. At P18, a significant Holm-Sidak’s multiple comparison post hoc test was observed between Long and Short (p = 0.0002) groups, and a trend level effect between Long and Equinox (p = 0.0886) photoperiod conditions. For DA content, using a two-way ANOVA, we found a significant main effect of photoperiod (p < 0.0001; F (2, 102) = 12.97) and a significant main effect of age (p = 0.0058; F (2, 102) = 5.422) **(Figure 3C)**. Holm-Sidak’s multiple comparison post hoc tests revealed a trend level effect between Long and Short photoperiods at P18 (p = 0.0501), and at P35 significant differences were found between Long and Short (p = 0.0043), and Equinox and Short (p = 0.0493) groups. For DOPAC content, using a two-way ANOVA, we observed a significant main effect of photoperiod (p = 0.0005; F (2, 102) = 8.133), and a significant main effect of age (p < 0.0001; F (2, 102) = 17.38) **(Figure 3D)**. At P35, a significant Holm-Sidak’s multiple comparison post hoc test was found between Long and Short photoperiods (p = 0.0089). Lastly, we measured norepinephrine and epinephrine values as well. For norepinephrine content, using a two-way ANOVA, we found a non-significant main effect of photoperiod (p = 0.1627; F (2, 102) = 1.849), a significant main effect of age (p < 0.0001; F (2, 102) = 287.9), and a significant interaction effect (p = 0.0124; F (4, 102) = 3.369) **(Supplementary Figure 3)**. Holm’s-Sidak’s multiple comparison post hoc tests revealed a significant difference between Short and Long photoperiods at P8 (p = 0.0245). For midbrain epinephrine content, values across age and group were so low that they were undetectable. Overall, we observed that developmental Short photoperiod exposure results in decreased levels of midbrain serotonin and dopamine content along with their corresponding metabolites, with these differences manifesting at P18 and P35, representing early childhood and adolescence in the mouse, respectively^68^.

**Figure 3.**
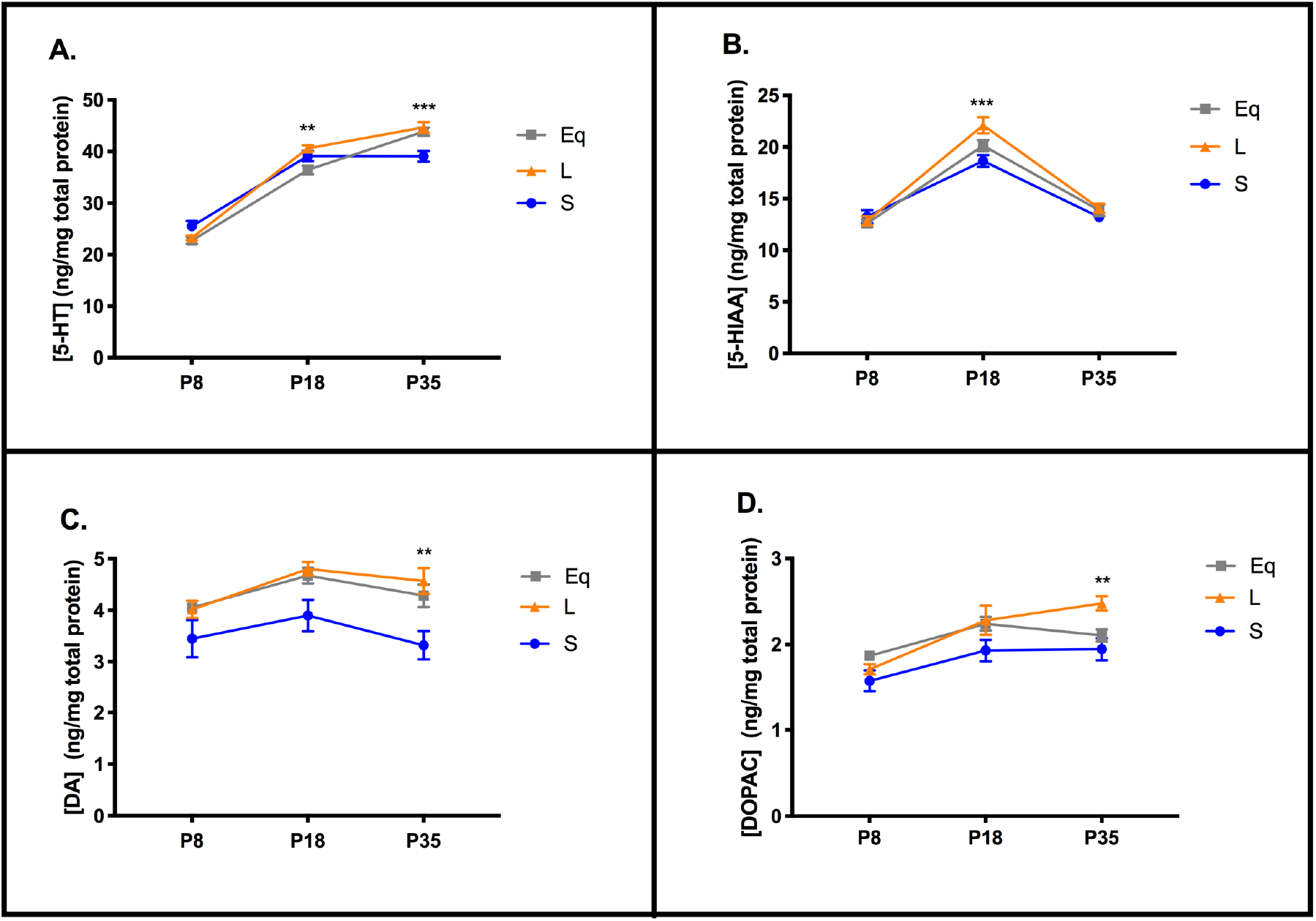
Short photoperiod reduced midbrain monoamine content at multiple stages of development. **A)** 5-HT content, **B)** 5-HIAA levels, **C)** DA content, and **D)** DOPAC levels in the midbrain. Mice developed under either Equinox (Eq), Long (L), or Short (S) photoperiods and monoamine levels were measured at three time points: P8, P18, and P35. The significant levels were as follows: (** = p < 0.01, *** = p < 0.001). The significance levels in A) are shown for Long (L) to Equinox (Eq) at P18, and for Long (S) to Short (S) comparisons at P35. In addition, a significant Holm-Sidak’s post hoc test revealed differences between Equinox and Short (p = 0.0014) groups for **A)** 5-HT content at P35. The significance levels in **B)** are shown for Long (S) to Short (S) comparisons at P18. The significance levels in **C)** and **D)** are shown for Long (L) to Short (S) comparisons at P35. Lastly, a significant Holm-Sidak’s post hoc test revealed differences between Equinox and Short (p = 0.0493) groups for **C)** DA content at P35.

### Midbrain Serotonergic Gene Expression in Early Adulthood is Significantly Reduced in Long Summer-like Photoperiod Mice

Based on the highest incidence of depression occurring during early adulthood^2^, we investigated the effects of photoperiod on expression levels of key 5-HT genes in the midbrain at P50, representing early adulthood in the mouse^71^. Quantitative RTPCR at P50 was used to evaluate the expression of serotonergic genes of interest (*Tph2, SERT* and *Pet-1*) for mice developed under control Equinox, Long summer-like, and Short winter-like photoperiod conditions from E0 **(Figure 4)**. Using a one-way ANOVA, a significant main effect of photoperiod was found for *Tph2* expression (p = 0.0118; F (2, 15) = 6.063) **(Figure 5A)**. In addition, a significant Holm-Sidak’s multiple comparison post hoc test was observed between Equinox and Long conditions (p = 0.0123) and a trend level effect was found comparing Short and Long photoperiods (p = 0.0574). Also, using a one-way ANOVA, a significant main effect of photoperiod was found for *SERT* expression (p = 0.0014; F (2, 15) = 10.56) with significant Holm-Sidak’s multiple comparison effects being observed between Equinox and Long photoperiod conditions (p = 0.0019), and between Short and Long photoperiods (p = 0.0058) **(Figure 5B)**. Lastly, no significant main effect of photoperiod was found for *Pet-1* expression (p = 0.4921; F (2, 15) = 0.7437) **(Figure 5C)**. By utilizing quantitative RTCPR in the midbrain during early adulthood, it was observed that *Tph2* and *SERT* expression levels are reduced under Long summer-like conditions compared to both Equinox control and Short winter-like photoperiodic conditions.

**Figure 4.**
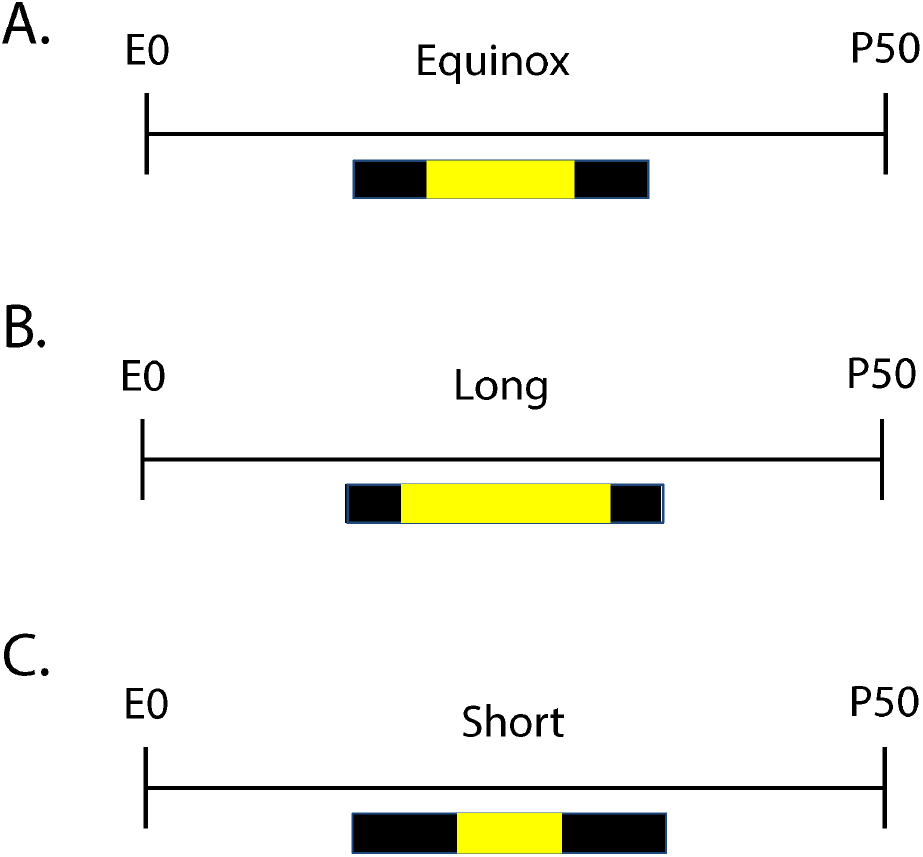
Schematic of developmental photoperiod paradigm. For early adulthood <u>midbrain RTPCR 5-HT gene experiments</u> animals developed under either an **A)** Equinox, **B)** Long, or **C)** Short photoperiod. For early adulthood DRN RNAScope <u>experiments animals</u> developed under either a **B)** Long or **C)** Short photoperiod. Animals developed under Equinox LD 12:12, Long LD 16:8, or Short LD 8:16 photoperiods from embryonic day 0 (E0) to postnatal day 50 (P50).

**Figure 5.**
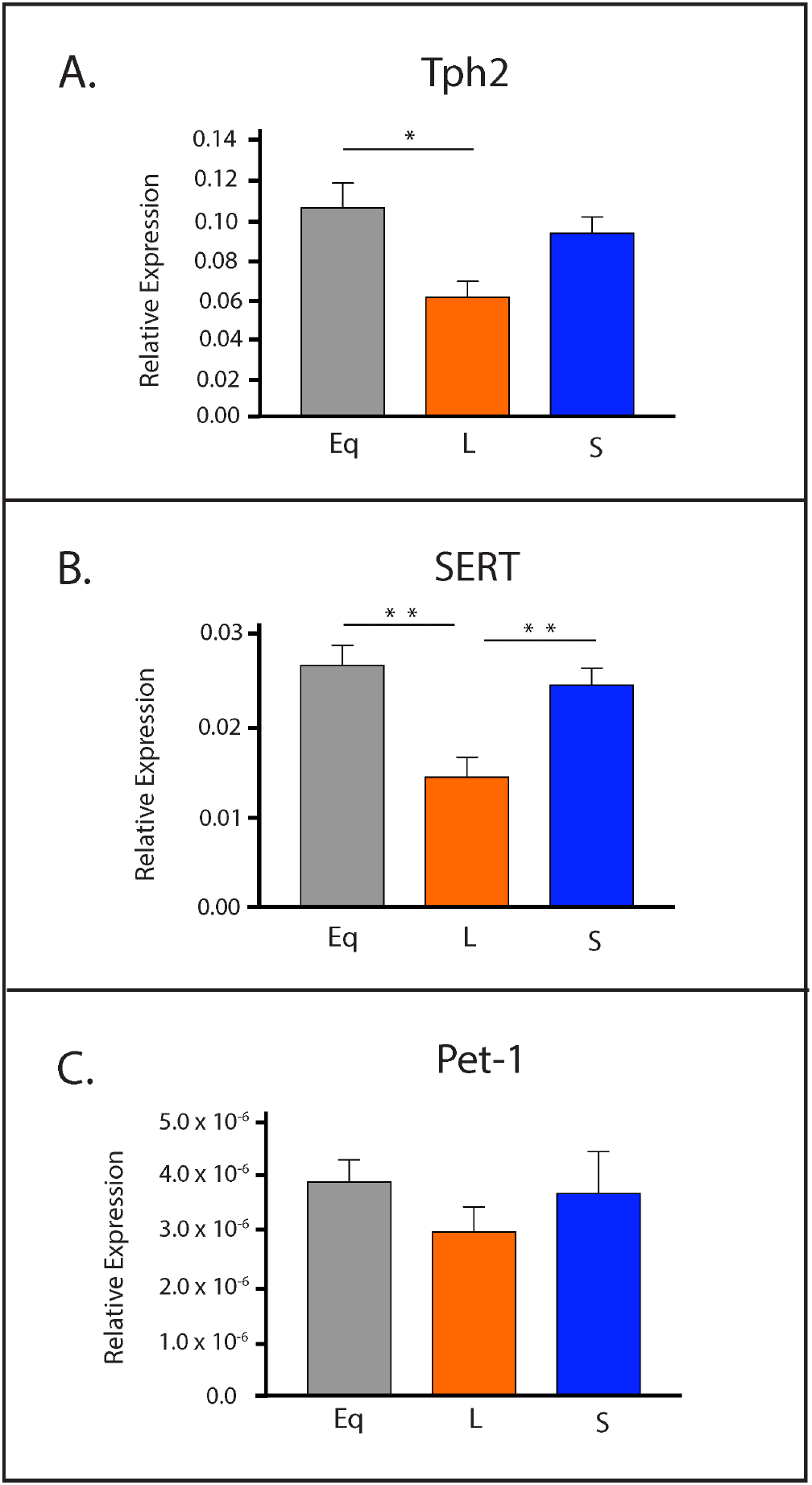
Long photoperiodic conditions resulted in significant decreases in expression levels of key 5-HT genes in the midbrain during early adulthood. **A)** *Tph2* expression, **B)** *SERT* expression, and **C)** *Pet-1* expression. Mice developed under either Equinox (Eq), Long (L), or Short (S) photoperiods and 5-HT gene expression levels were measured at P50. The significant levels are as follows: (* = p < 0.05, ** = p < 0.01). Note a trend level effect when comparing Short (S) and Long (L) photoperiods for *Tph2* expression **A)** (p = 0.0574). Reference gene for relative expression was *Hprt*.

### RNAScope Reveals Reduced Adult *Tph2* Expression Levels in Mice Developed Under Long Summer-like Photoperiod Conditions in the DRN

In addition, we wanted to investigate the role of developmental photoperiod on adult gene expression levels in the main hub for serotonin synthesis in the brain, the dorsal raphe nucleus (DRN) by utilizing RNAScope methods. Mice developed at E0 under either Long or Short photoperiods and were evaluated at P50 **(Figure 4B-C)**. Representative images of the DRN were obtained for animals that developed under Long summer-like or Short winter-like photoperiods with fluorescent dyes Alexa488 staining *Tph2* in green, Atto550 staining *SERT* in red, Atto655 staining *Pet-1* in blue, and the three images were then merged to evaluate the gene expression overlap for all cells within the DRN **(Figure 6)**. Paired t tests revealed significant differences between Short and Long photoperiods for integrated density of cell fluorescence (p = 0.0137) **(Figure 7A)**, a trend level effect for integrated density of DRN (i.e. ROI) fluorescence (p = 0.0616) **(Figure 7B)**, significant differences in mean cell fluorescence (p = 0.0362) **(Figure 7C)**, and in ROI mean fluorescence (p = 0.0252) **(Figure 7D)**. Paired t tests revealed no significant differences in *SERT* expression between Short and Long photoperiods for integrated density of cell fluorescence (p = 0.2515), integrated density of ROI fluorescence (p = 0.1331), mean cell fluorescence (p = 0.1379), or in ROI mean fluorescence (p = 0.1269) **(Supplementary Figure 4)**. Interestingly, while no significant differences were observed between the groups, *SERT* expression was elevated in all four measures for Short compared to Long photoperiod conditions **(Supplementary Figure 4)**. In addition, no significant differences were observed in *Pet-1* expression between photoperiods when evaluating integrated density of cell fluorescence (p = 0.2822), integrated density of ROI fluorescence (p = 0.2822), mean cell fluorescence (p = 0.1270), or ROI mean fluorescence (p = 0.1270) **(Supplementary Figure 5)**. Lastly, no significant differences in total cell number were observed between Short and Long photoperiods for *Tph2* (p = 0.9872), *SERT*(p = 0.3238), or *Pet-1* expression (p > 0.9999) **(Supplementary Figure 6)**. In addition, the number of cells appeared to be comparable across the evaluated serotonergic genes: *Tph2* (Short – 282.3 ± 18.4, Long – 282.7 ± 9.996), *SERT*(Short – 288 ± 14.1, Long – 269.3 ± 8.724), and *Pet-1* (Short – 267.8 ± 11.17, Long – 267.8 ± 10.03). By utilizing RNAScope methods targeting the DRN during early adulthood, we found *Tph2* expression to be significantly reduced for Long compared to Short photoperiod mice using four different measures, and *SERT*, *Pet-1* expression, and the total cell numbers were not significantly different between the two photoperiod conditions.

**Figure 6.**
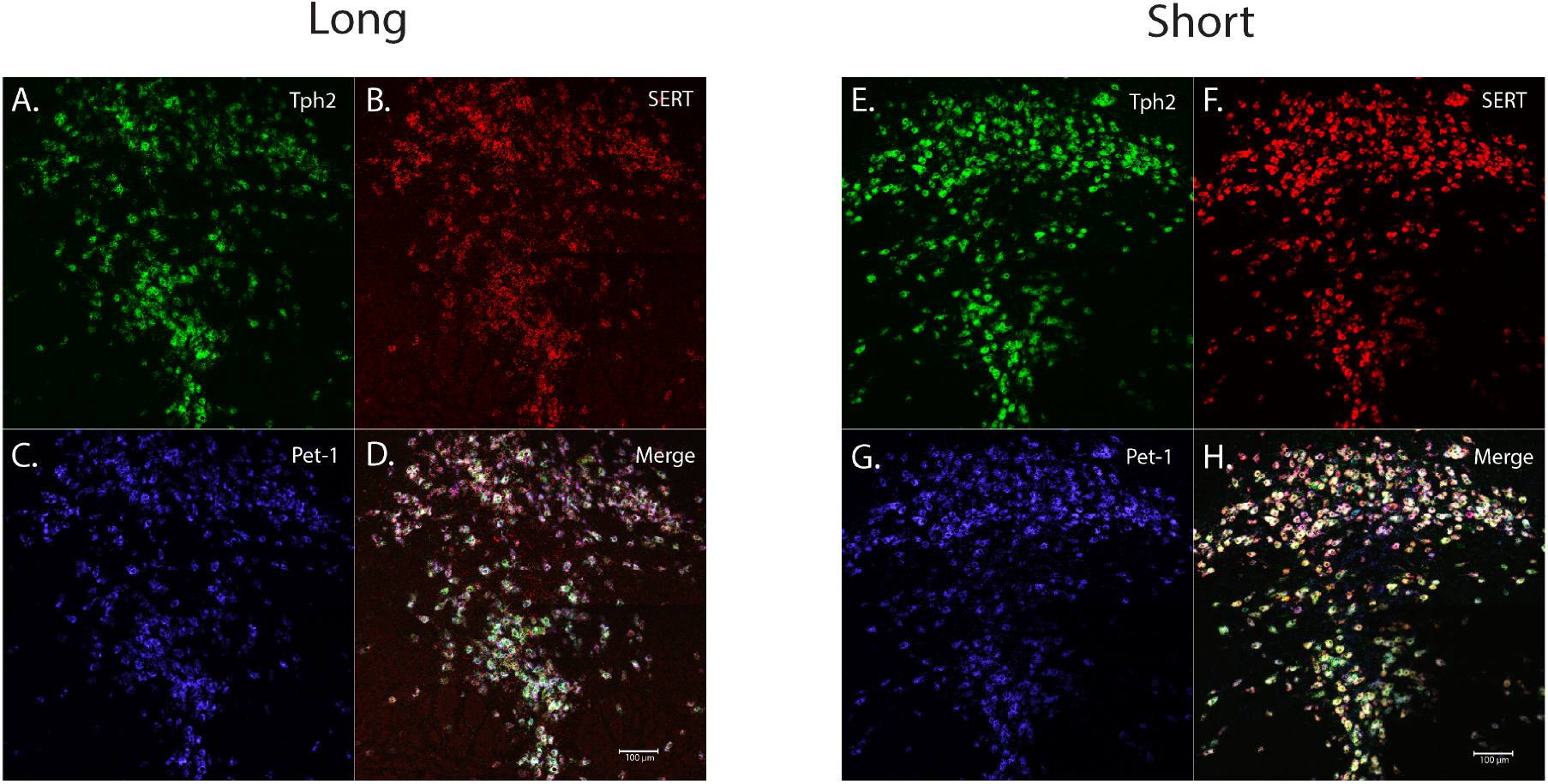
RNAScope revealed significant differences in adult gene expression of relevant 5-HT genes in the DRN due to developmental photoperiod. Representative image for an animal that developed under a Long photoperiod and measured at P50: **A)** *Tph2* expression is stained in green and tagged with Alexa488, **B)** *SERT* expression is stained in red and tagged with Atto550, **C)** *Pet-1* expression is stained in blue and tagged with Atto647, and **D)** is the overlay of all three channels. Representative image for an animal that developed under a Short photoperiod and measured at P50: **E)** *Tph2* expression is stained in green and tagged with Alexa488, **F)** *SERT* expression is stained in red and tagged with Atto550, **G)** *Pet-1* expression is stained in blue and tagged with Atto647, and **H)** is the overlay of all three channels. Note that qualitatively, fluorescent levels are brighter under Short compared to Long photoperiod conditions. Scale bar was set at 100μm.

**Figure 7.**
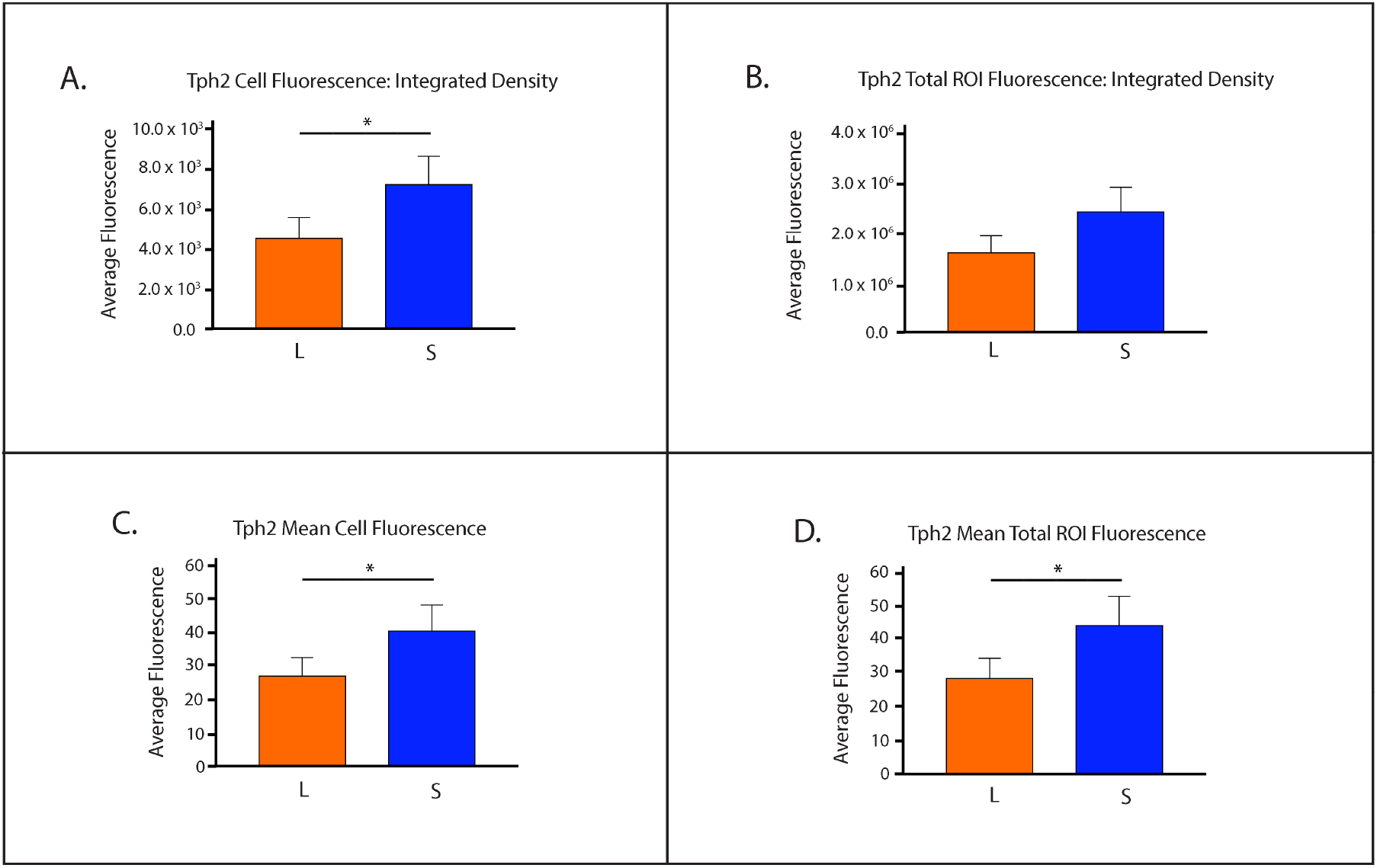
*Tph2* expression levels were significantly decreased in the DRN of mice developed under Long photoperiod conditions in early adulthood. **A)** Integrated density of cell fluorescence, **B)** Integrated density of ROI fluorescence, **C)** Quantification of mean cell fluorescence, and **D)** Quantification of ROI mean fluorescence. Mice developed under either Long (L) or Short (S) photoperiods and RNAScope experiments occurred at P50. The significant levels are as follows: (* = p < 0.05). Note a trend level effect when comparing Short and Long photoperiods for Integrated density of ROI fluorescence **B)** (p = 0.0616).

## Discussion

In this study we evaluated the effects of photoperiod during key periods of postnatal development on gene expression levels and midbrain monoamine content in the dopaminergic and serotonergic systems. We found that gene expression levels of *TH*, the rate-limiting enzyme for dopamine synthesis, were elevated in Short winter-like photoperiod mice. This difference was specifically observed in P8 animals, representing perinatal development^68^, and this increase in *TH* expression gradually decreased, reaching trend level differences by P18, and eventually normalizing by P35. When evaluating serotonergic gene expression (i.e. *Tph2, SERT, Pet-1),* we observed that levels were not altered by photoperiod throughout development. In addition, we observed that midbrain serotonin (5-HT) and dopamine (DA) tissue content along with their corresponding metabolites (5-HIAA and DOPAC), were significantly reduced in mice developed under Short winter-like compared to Long summer-like photoperiods. This suggests that Short winter-like photoperiod exposure can decrease not only dopamine and serotonin content levels, but can reduce monoamine utilization, as observed via elevated metabolite concentrations. Interestingly, these differences manifested at either P18 and/or P35, representing early childhood and adolescence in the mouse^68^. Overall, when investigating through development, midbrain *TH* expression levels were elevated earlier in development, by P8, in Short winter-like mice, normalizing by P35 and midbrain monoamine content was significantly reduced in Short photoperiod animals by P18 and P35.

We also focused on early adulthood, as this is a consistent time period associated with mood disorders^7^, to assess the expression of key serotonergic genes in the midbrain. Interestingly, at P50 we found that *Tph2* and *SERT* expression levels were reduced for Long photoperiod animals compared to both Equinox and Short photoperiod conditions, while no differences were observed for *Pet-1* expression. We then utilized RNAScope methods to specifically target the known key brain region for serotonin synthesis and neuron development (i.e. the DRN). We found that *Tph2* expression was again significantly reduced for mice that developed under Long summer-like compared to Short winter-like photoperiods for integrated density of cell fluorescence, integrated density of ROI fluorescence, mean cell fluorescence, and ROI mean fluorescence. *SERT* and *Pet-1* expression were comparable between photoperiods, however *SERT* expression levels were consistently reduced in Long photoperiod animals for all four fluorescence measures. Lastly, there were no significant differences in total cell number observed between photoperiods or when evaluating these genes of interest. Importantly, this demonstrated that the current findings are not due to simply an overall increase in cell number, but rather consistently reduced gene expression levels in DRN cells for Long photoperiod animals. Therefore, it appears that with two different measures and locations, quantitative RTPCR in the midbrain and RNAScope in the DRN, *Tph2* and possibly *SERT* expression are reduced in animals raised under Long summer-like compared to Short winter-like photoperiods by early adulthood (P50), which could not be explained by potential differences in total cell numbers.

We recently demonstrated that Long developmental photoperiod can program DRN 5-HT neuronal firing rate *prenatally* in mice, whereas Long photoperiods can modulate monoamine content and the resulting affective behaviors during *postnatal* development^34^ Therefore, we proposed a double hit hypothesis in which photoperiodic programming of the serotonin system may occur sequentially, impacting DRN 5-HT neurons prenatally, and then modulating monoamine signaling responsible for the underlying circuitry and resultant affective behaviors during specific periods of postnatal development^34^ While we have previously focused on the DRN, in the current study we aimed to determine these sensitive postnatal periods in the midbrain, which encompasses aspects of both the serotonergic and dopaminergic systems.

Studies have identified sensitive periods in which monoamine content, receptors, and the respective transporters of the dopamine and serotonin systems develop and the role these time windows may have in the development of affective disorders^74^. In the current study we observed similar effects, as found in early adulthood, for monoamine signaling of 5-HT, DA, and their corresponding metabolites, such that monoamine content was significantly reduced in mice developed under Short winter-like compared to Long summer-like photoperiods, however these effects arose by P18 or P35. These time periods mirror windows that are critical for the development of key aspects of the dopamine and serotonin systems^74^. In addition, this has intriguing implications as these periods represent childhood and early adolescence in the mouse^68^, respectively, and have been associated with mood disorders in humans^6^. Thus, we may have identified sensitive periods of postnatal development, which may be vulnerable to the effects of an environmental factor such as photoperiod, which could then impact the underlying circuitry responsible for the resultant affective behavior.

While previous studies have found intriguing effects of photoperiod in the dopaminergic system due to photoperiod^29–31^, no study has investigated photoperiodic effects due to *developmental* photoperiod, across multiple stages of development, or in mice that are melatonin competent, such as the C3Hf^+/+^ strain. In the current study we observed reduced levels of DA and DOPAC for Short winter-like photoperiod mice at P35, thus based on this data and prior work, we hypothesized that the expression of a key dopamine gene, *TH*, would be decreased for animals raised under Short photoperiodic conditions as well. Interestingly, we found that midbrain *TH* expression was comparable across photoperiods at P35, but that mice developed under Short winter-like photoperiods demonstrated a significant increase in midbrain *TH* expression compared to Long photoperiod animals at P8. While transcriptional regulation of *TH* is important, there are multiple post-translational mechanisms of *TH* regulation as well including phosphorylation, degradation, and protein binding^75^, which may explain these differences.

In addition, we observed age-dependent effects on midbrain *Tph2,* with levels highest at P8 and then declining, while *SERT* expression in the midbrain showed no main effects of age or photoperiod from P8 through P35. However, by P50, there were clear photoperiod-dependent effects on the midbrain expression of *Tph2* and *SERT* although not on the 5-HT neuron-specific transcription factor *Pet-1*, with *Tph2* and *SERT* being reduced in Long photoperiods. The tendency for *Tph2* and *SERT* to be reduced in Long photoperiods was also observed in RNAScope assays targeting DRN 5-HT neurons specifically. Thus, these key serotonin signaling genes are consistently found to be down-regulated in Long photoperiods by P50, conditions when serotonin content has been found to be elevated in the dorsal raphe nucleus^15^. This suggests that there may be potential compensatory mechanisms established for both dopaminergic and serotonergic gene expression levels in animals raised under Long summer-like photoperiods.

It is plausible that the previously observed increase in firing rate of 5-HT neurons in the DRN may be driving the increased levels in monoamine content in the midbrain found in animals raised under Long photoperiods. Serotonin and norepinephrine midbrain content has been shown to be significantly increased in Long photoperiod mice by P50^15,34^, yet in the current study *Tph2* and *SERT* expression are significantly decreased in Long photoperiod animals at this age. Based on the significant increase in monoamine content observed in Long photoperiod animals, 5-HT gene expression levels in both the midbrain and the hub of 5-HT synthesis, the DRN, may be down regulated to compensate for these elevated monoamine tissue concentrations. Therefore, if the serotonergic system is being driven excessively under Long photoperiod conditions it may be possible that compensatory gene expression mechanisms may be in place to slow or down regulate these elevated levels. Along the same line, Short winter-like photoperiods may up-regulate gene expression levels to compensate for the reduced midbrain content observed in the dopamine and serotonin systems. While no study has previously evaluated dopamine gene expression levels due to developmental photoperiod in melatonin competent mice, we found *TH* levels dramatically increase during perinatal development (P8), normalizing by P35 or early adolescence in the mouse. Studies have shown that the perinatal window can be a sensitive period for the dopamine system and therefore this up-regulation of gene expression may result in dramatic changes to the underlying DA neurons, circuits, and behaviors^74^, which we at least observed peripherally with DA and DOPAC content later in the course of development. These findings may suggest that depending on the time period, gene expression can be modulated differentially by photoperiod depending on the neurotransmitter system evaluated. Regardless, it is clear that gene expression is up-regulated in Short winter-like mice in the dopamine system, at P8, and down-regulated in Long summer-like mice in the serotonin system, at P50, which could result in lasting changes to the underlying circuitry and related behaviors.

The effects of photoperiod during development on firing rate in DRN 5-HT neurons, midbrain monoamine concentrations, behaviors related to anxiety and depression, and now gene expression changes have been investigated during early adulthood in the rodent model^15,33,37,38^. However, it would be highly relevant to assess potential changes in neural activity, signaling, and the associated behaviors at multiple time points during development to more fully understand the effects that environmental factors such as photoperiod have on the dopamine and serotonin systems. To this point, *Tph2, SERT,* and *Pet-1* expression should be evaluated at multiple developmental sensitive periods in the DRN to examine the overall trajectory that photoperiodic effects may have on serotonergic gene expression and gain further insight into if these changes are due to compensatory mechanisms as hypothesized above. In addition, there is a known population of dopaminergic cells in the DRN^76^ and DRN dopamine cells have been shown to play an role in arousal and are activated by salient stimuli^77^. Therefore, future studies should evaluate the expression of *TH* and *DAT* in both the midbrain and DRN during multiple stages of development including early adulthood. Also, studies have shown that serotonin neuronal synthesis, projections, and the overall development of the serotonergic system can occur as early as between embryonic days 12-14 in rodents^59,63,78–83^. This suggests that photoperiodic exposure during perinatal or prenatal time points may have lasting effects on serotonin gene expression. In this study, we evaluated photoperiodic effects during the course of development, and identified time periods in which monoamine and gene expression changes occurred. However, this does not unequivocally identify these time windows as sensitive periods. To address this, future studies may switch animals to different photoperiods during either prenatal or perinatal development to determine if these effects on gene expression are driven by developmental or proximal photoperiods.

In addition to these basic science findings, this study has potentially significant clinical implications. The dopaminergic and serotonergic systems have been implicated in various neurodevelopmental disorders such as major depressive disorder, anxiety, and autism spectrum disorder^39,40,84–89^. With preclinical models it has been shown that modulation of serotonin and dopamine during key developmental time points can vastly alter neuronal firing, circuit formation, and the associated behaviors^90–95^ As we are beginning to identify the underlying mechanisms and the sensitive postnatal developmental periods impacted by the duration of daylight or photoperiod in the mouse model, this may be clinically relevant, as there is evidence to suggest that light therapy may be effective in treating children and adolescents with seasonal affective disorder^99^ and major depression^100,101^.

Overall, it was found that mice developed under Short winter-like photoperiods demonstrate reduced midbrain serotonin, dopamine, 5-HIAA, and DOPAC content compared to Long summer-like photoperiod mice, with these differences manifesting by P18 and P35. This work follows similar results observed in multiple rodent species evaluated in early adulthood and suggests that these time periods, representing childhood and early adolescence in the mouse, may represent vulnerable periods in postnatal development. Interestingly, we observed that midbrain *TH* levels were significantly increased for Short photoperiod animals during perinatal development, at P8, and normalized by P35. In early adulthood, P50, we showed that animals raised under a Long photoperiod demonstrate decreased expression levels of *Tph2* and *SERT*compared to mice that developed under either a Short or Equinox photoperiod. We observed similar photoperiodic effects using both quantitative RTPCR in the midbrain and RNAScope in the DRN, which were not driven by potential differences in cell number. Based on prior results, we hypothesize that there is an up-regulation of genes relevant to the dopaminergic (*TH*) system in Short photoperiod mice as observed during perinatal development, and a down-regulation of genes relevant to the serotonergic (*Tph2* and *SERT*) system in early adulthood, (P50). Interestingly, these findings may indicate that depending on the sensitive period, gene expression is modulated differentially depending on the neurotransmitter system, potentially resulting in enduring changes to the underlying circuitry and related behaviors later in life. Thus, investigating the interactions between photoperiod, monoamine signaling, and gene expression levels in the dopaminergic and serotonergic systems during the course of development may provide novel insights into the etiology, underlying mechanisms, and potential therapeutic targets for mood disorders.

## Acknowledgments

This research was supported by National Institutes of Health Grants NIH R01 MH108562 to DGM, 5T32MH018921-24: Development of Psychopathology: From Brain and Behavioral Science to Intervention (JKS), 1P50MH096972 from the Vanderbilt University Silvio O. Conte Center, the Vanderbilt University Conte Center Pilot Project (JKS), Joel G. Hardman and Mary K. Parr Endowed Chair in Pharmacology (RE), R01 DK119508 (RE), and the Simms/Mann Chair in Neurodevelopmental Neurogenetics (PL). Experiments using the Zeiss LSM510 Confocal microscope were performed in part through the use of the VUMC Cell Imaging Shared Resource (supported by NIH grants CA68485, DK20593, DK58404, DK59637, and EY08126). We would also like to thank Sean Schaffer for his help and advice on this project. We would also like to thank Maia Weisenhaus for her help with confocal microscope imaging for the RNAScope aspect of this project. The Vanderbilt Neurochemistry Core is supported by the EKS NICHD of the NIH under award U54HD083211.

## Author Contributions

JS, TM, RE, and DM wrote the manuscript. JS collected the samples and analyzed the data for the midbrain RTCPR experiments, JS imaged, quantified, and analyzed the data for the RNAScope experiments, CJ and NH performed the DRN RTPCR experiments, TM performed the midbrain RTPCR experiments, NH collected the samples and PW performed the RNAScope staining for the RNAScope experiments. JS, TM, CJ, NH, PW, RE, PL, and DM all discussed aspects of the experimental designs, procedures, and overall project ideas.

## References

1 (WHO), W. H. O. Depression. (2019).

2 (NIMH), N. I. o. M. H. Major Depression Among Adults. (2019).

3 (NIMH), N. I. o. M. H. Major Depression Among Adolescents. (2019).

4 (CDC), C. f D. C. a. P. Data and Statistics on Children’s Mental Health, <https://www.cdc.gov/childrensmentalhealth/data.html> (2019).

5 Hammen, C., Garber, J. & Ingram, R. E. Vulnerability to depression across the lifespan. Vulnerability to psychopathology: Risk across the lifespan, 258–267 (2001).

6 Bhatia, S. K. & Bhatia, S. C. Childhood and adolescent depression. Depression 100, 53 (2007).

7 Rohde, P., Lewinsohn, P. M., Klein, D. N., Seeley, J. R. & Gau, J. M. Key characteristics of major depressive disorder occurring in childhood, adolescence, emerging adulthood, and adulthood. Clinical Psychological Science 1, 41–53 (2013).

8 Nestler, E. J. et al. Neurobiology of depression. Neuron 34, 13–25 (2002).

9 Andersen, S. L. Exposure to early adversity: points of cross-species translation that can lead to improved understanding of depression. Development and psychopathology 27, 477–491 (2015).

10 Modai, I., Kikinzon, L. & Valevski, A. Environmental factors and admission rates in patients with major psychiatric disorders. Chronobiology international 11, 196–199 (1994).

11 Castrogiovanni, P., Iapichino, S., Pacchierotti, C. & Pieraccini, F. Season of birth in psychiatry. Neuropsychobiology 37, 175–181 (1998).

12 Tonetti, L., Fabbri, M., Martoni, M. & Natale, V. Season of birth and mood seasonality in late childhood and adolescence. Psychiatry research 195, 66–68 (2012).

13 Park, D. H., Kripke, D. F. & Cole, R. J. More prominent reactivity in mood than activity and sleep induced by differential light exposure due to seasonal and local differences. Chronobiology international 24, 905–920 (2007).

14 Qin, D. et al. The first observation of seasonal affective disorder symptoms in Rhesus macaque. Behavioural brain research 292, 463–469 (2015).

15 Green, N. H., Jackson, C. R., Iwamoto, H., Tackenberg, M. C. & McMahon, D. G. Photoperiod programs dorsal raphe serotonergic neurons and affective behaviors. Current Biology 25, 1389–1394 (2015).

16 Xu, L.-Z. et al. Short photoperiod condition increases susceptibility to stress in adolescent male rats. Behavioural brain research 300, 38–44 (2016).

17 Otsuka, T. et al. Photoperiodic responses of depression-like behavior, the brain serotonergic system, and peripheral metabolism in laboratory mice. Psychoneuroendocrinology 40, 37–47 (2014).

18 Disanto, G. et al. Seasonal distribution of psychiatric births in England. PloS one 7, e34866 (2012).

19 Torrey, E. F., Miller, J., Rawlings, R. & Yolken, R. H. Seasonality of births in schizophrenia and bipolar disorder: a review of the literature. Schizophrenia research 28, 1–38 (1997).

20 Zhang, L. et al. A PERIOD3 variant causes a circadian phenotype and is associated with a seasonal mood trait. Proceedings of the National Academy of Sciences 113, E1536–E1544 (2016).

21 Foster, R. G. & Roenneberg, T. Human responses to the geophysical daily, annual and lunar cycles. Current biology 18, R784–R794 (2008).

22 Chotai, J. & Adolfsson, R. Converging evidence suggests that monoamine neurotransmitter turnover in human adults is associated with their season of birth. European Archives of Psychiatry and Clinical Neuroscience 252, 130–134 (2002).

23 Lambert, G., Reid, C., Kaye, D., Jennings, G. & Esler, M. Effect of sunlight and season on serotonin turnover in the brain. The Lancet 360, 1840–1842 (2002).

24 Chotai, J., Serretti, A., Lattuada, E., Lorenzi, C. & Lilli, R. Gene-environment interaction in psychiatric disorders as indicated by season of birth variations in tryptophan hydroxylase (TPH), serotonin transporter (5-HTTLPR) and dopamine receptor (DRD4) gene polymorphisms. Psychiatry research 119, 99–111 (2003).

25 Devore, E. E., Chang, S.-C., Okereke, O. I., McMahon, D. G. & Schernhammer, E. S. Photoperiod during maternal pregnancy and lifetime depression in offspring. Journal of psychiatric research 104, 169–175 (2018).

26 Ciarleglio, C., Resuehr, H. & McMahon, D. Interactions of the serotonin and circadian systems: nature and nurture in rhythms and blues. Neuroscience 197, 8–16 (2011).

27 Caspi, A. et al. Influence of life stress on depression: moderation by a polymorphism in the 5-HTT gene. Science 301, 386–389 (2003).

28 Spindelegger, C. et al. Light-dependent alteration of serotonin-1A receptor binding in cortical and subcortical limbic regions in the human brain. The World Journal of Biological Psychiatry 13, 413–422 (2012).

29 Aumann, T. D. et al. Differences in number of midbrain dopamine neurons associated with summer and winter photoperiods in humans. PloS one 11, e0158847 (2016).

30 Young, J. W. et al. Mice with reduced DAT levels recreate seasonal-induced switching between states in bipolar disorder. Neuropsychopharmacology 43, 1721 (2018).

31 Dulcis, D., Jamshidi, P., Leutgeb, S. & Spitzer, N. C. Neurotransmitter switching in the adult brain regulates behavior. Science 340, 449–453 (2013).

32 Dominguez – López, S., Howell, R. D., López – Canúl, M. G., Leyton, M. & Gobbi, G. Electrophysiological characterization of dopamine neuronal activity in the ventral tegmental area across the light-dark cycle. Synapse 68, 454–467 (2014).

33 Goda, R. et al. Serotonin levels in the dorsal raphe nuclei of both chipmunks and mice are enhanced by long photoperiod, but brain dopamine level response to photoperiod is species-specific. Neuroscience letters 593, 95–100 (2015).

34 Siemann, J. K., Green, N. H., Reddy, N. & McMahon, D. G. Sequential photoperiodic programing of serotonin neurons, signaling and behaviors during prenatal and postnatal development. Frontiers in neuroscience 13, 459 (2019).

35 Itzhacki, J., Clesse, D., Goumon, Y., Van Someren, E. J. & Mendoza, J. Light rescues circadian behavior and brain dopamine abnormalities in diurnal rodents exposed to a winter-like photoperiod. Brain Structure and Function 223, 2641–2652 (2018).

36 Krivisky, K., Ashkenazy, T., Kronfeld-Schor, N. & Einat, H. Antidepressants reverse short-photoperiod-induced, forced swim test depression-like behavior in the diurnal fat sand rat: further support for the utilization of diurnal rodents for modeling affective disorders. Neuropsychobiology 63, 191–196 (2011).

37 Prendergast, B. J. & Nelson, R. J. Affective responses to changes in day length in Siberian hamsters (Phodopus sungorus). Psychoneuroendocrinology 30, 438–452 (2005).

38 Pyter, L. M. & Nelson, R. J. Enduring effects of photoperiod on affective behaviors in Siberian hamsters (Phodopus sungorus). Behavioral neuroscience 120, 125–134 (2006).

39 Nestler, E. J. & Carlezon Jr, W. A. The mesolimbic dopamine reward circuit in depression. Biological psychiatry 59, 1151–1159 (2006).

40 Dunlop, B. W. & Nemeroff, C. B. The role of dopamine in the pathophysiology of depression. Archives of general psychiatry 64, 327–337 (2007).

41 Chaudhury, D. et al. Rapid regulation of depression-related behaviours by control of midbrain dopamine neurons. Nature 493, 532 (2013).

42 Friedman, A. K. et al. Enhancing depression mechanisms in midbrain dopamine neurons achieves homeostatic resilience. Science 344, 313–319 (2014).

43 Serretti, A. et al. Tyrosine hydroxylase gene associated with depressive symptomatology in mood disorder. American journal of medical genetics 81, 127–130 (1998).

44 Waløen, K., Kleppe, R., Martinez, A. & Haavik, J. Tyrosine and tryptophan hydroxylases as therapeutic targets in human disease. Expert opinion on therapeutic targets 21, 167–180 (2017).

45 Zhu, M.-Y. et al. Elevated levels of tyrosine hydroxylase in the locus coeruleus in major depression. Biological psychiatry 46, 1275–1286 (1999).

46 Gaspar, P., Cases, O. & Maroteaux, L. The developmental role of serotonin: news from mouse molecular genetics. Nature Reviews Neuroscience 4, 1002–1012 (2003).

47 Gross, C. et al. Serotonin1A receptor acts during development to establish normal anxiety-like behaviour in the adult. Nature 416, 396–400 (2002).

48 Holmes, A., Murphy, D. L. & Crawley, J. N. Abnormal behavioral phenotypes of serotonin transporter knockout mice: parallels with human anxiety and depression. Biological psychiatry 54, 953–959 (2003).

49 Bennett, A. J. et al. Early experience and serotonin transporter gene variation interact to influence primate CNS function. Molecular psychiatry 7, 118 (2002).

50 Uher, R. & McGuffin, P. (Nature Publishing Group, 2008).

51 Eley, T. C. et al. Gene-environment interaction analysis of serotonin system markers with adolescent depression. Molecular psychiatry 9, 908–915 (2004).

52 Bennett-Clarke, C. A., Leslie, M. J., Chiaia, N. L. & Rhoades, R. W. Serotonin 1B receptors in the developing somatosensory and visual cortices are located on thalamocortical axons. Proceedings of the National Academy of Sciences 90, 153–157 (1993).

53 Bonnin, A. et al. A transient placental source of serotonin for the fetal forebrain. Nature 472, 347–350 (2011).

54 Bonnin, A. & Levitt, P. Fetal, maternal, and placental sources of serotonin and new implications for developmental programming of the brain. Neuroscience 197, 1–7 (2011).

55 Cases, O. et al. Aggressive behavior and altered amounts of brain serotonin and norepinephrine in mice lacking MAOA. Science (New York, NY) 268, 1763 (1995).

56 Cases, O. et al. Lack of barrels in the somatosensory cortex of monoamine oxidase A-deficient mice: role of a serotonin excess during the critical period. Neuron 16, 297–307 (1996).

57 Esaki, T. et al. Developmental disruption of serotonin transporter function impairs cerebral responses to whisker stimulation in mice. Proc Natl Acad Sci U S A 102, 5582–5587 (2005).

58 Ul’yana, A. B., Bondar, N. P., Filipenko, M. L. & Kudryavtseva, N. N. Downregulation of serotonergic gene expression in the Raphe nuclei of the midbrain under chronic social defeat stress in male mice. Molecular neurobiology 48, 13–21 (2013).

59 Azmitia, E. C. Modern views on an ancient chemical: serotonin effects on cell proliferation, maturation, and apoptosis. Brain research bulletin 56, 413–424 (2001).

60 Kim, D.-K. et al. Altered serotonin synthesis, turnover and dynamic regulation in multiple brain regions of mice lacking the serotonin transporter. Neuropharmacology 49, 798–810 (2005).

61 Persico, A. M. et al. Reduced programmed cell death in brains of serotonin transporter knockout mice. Neuroreport 14, 341–344 (2003).

62 Deneris, E. S. & Wyler, S. C. Serotonergic transcriptional networks and potential importance to mental health. Nature neuroscience 15, 519–527 (2012).

63 Hendricks, T., Francis, N., Fyodorov, D. & Deneris, E. S. The ETS domain factor Pet-1 is an early and precise marker of central serotonin neurons and interacts with a conserved element in serotonergic genes. Journal of Neuroscience 19, 10348–10356 (1999).

64 Hendricks, T. J. et al. Pet-1 ETS gene plays a critical role in 5-HT neuron development and is required for normal anxiety-like and aggressive behavior. Neuron 37, 233–247 (2003).

65 Lukkes, J. L., Kopelman, J. M., Donner, N. C., Hale, M. W. & Lowry, C. A. Development× environment interactions control tph2 mRNA expression. Neuroscience 237, 139–150 (2013).

66 Oh, J.-e., Zupan, B., Gross, S. & Toth, M. Paradoxical anxiogenic response of juvenile mice to fluoxetine. Neuropsychopharmacology 34, 2197–2207 (2009).

67 Salichon, N. et al. Excessive activation of serotonin (5-HT) 1B receptors disrupts the formation of sensory maps in monoamine oxidase a and 5-ht transporter knock-out mice. Journal of Neuroscience 21, 884–896 (2001).

68 Semple, B. D., Blomgren, K., Gimlin, K., Ferriero, D. M. & Noble-Haeusslein, L. J. Brain development in rodents and humans: Identifying benchmarks of maturation and vulnerability to injury across species. Progress in neurobiology 106, 1–16 (2013).

69 Schmidt, S. Y. & Lolley, R. N. Cyclic-nucleotide phosphodiesterase: An early defect in inherited retinal degeneration of C3H mice. The Journal of cell biology 57, 117–123 (1973).

70 Ciarleglio, C. M., Axley, J. C., Strauss, B. R., Gamble, K. L. & McMahon, D. G. Perinatal photoperiod imprints the circadian clock. Nature neuroscience 14, 25–27 (2011).

71 Hefner, K. & Holmes, A. Ontogeny of fear-, anxiety-and depression-related behavior across adolescence in C57BL/6J mice. Behavioural brain research 176, 210–215 (2007).

72 Livak, K. J. & Schmittgen, T. D. Analysis of relative gene expression data using real-time quantitative PCR and the 2-ΔΔCT method. methods 25, 402–408 (2001).

73 Kirby, L., Pernar, L., Valentino, R. & Beck, S. Distinguishing characteristics of serotonin and non-serotonin-containing cells in the dorsal raphe nucleus: electrophysiological and immunohistochemical studies. Neuroscience 116, 669–683 (2003).

74 Suri, D., Teixeira, C. M., Cagliostro, M. K. C., Mahadevia, D. & Ansorge, M. S. Monoamine-sensitive developmental periods impacting adult emotional and cognitive behaviors. Neuropsychopharmacology 40, 88 (2015).

75 Daubner, S. C., Le, T. & Wang, S. Tyrosine hydroxylase and regulation of dopamine synthesis. Archives of biochemistry and biophysics 508, 1–12 (2011).

76 Stratford, T. R. & Wirtshafter, D. Ascending dopaminergic projections from the dorsal raphe nucleus in the rat. Brain research 511, 173–176 (1990).

77 Cho, J. R. et al. Dorsal raphe dopamine neurons modulate arousal and promote wakefulness by salient stimuli. Neuron 94, 1205–1219. e1208 (2017).

78 Aitken, A. R. & Törk, I. Early development of serotonin – containing neurons and pathways as seen in wholemount preparations of the fetal rat brain. Journal of Comparative neurology 274, 32–47 (1988).

79 Lauder, J. M. & Krebs, H. Serotonin as a differentiation signal in early neurogenesis. Developmental neuroscience 1, 15–30 (1978).

80 Wallace, J. A. & Lauder, J. M. Development of the serotonergic system in the rat embryo: an immunocytochemical study. Brain research bulletin 10, 459–479 (1983).

81 Lidov, H. G. & Molliver, M. E. Immunohistochemical study of the development of serotonergic neurons in the rat CNS. Brain research bulletin 9, 559–604 (1982).

82 Vitalis, T. & Parnavelas, J. G. The role of serotonin in early cortical development. Developmental neuroscience 25, 245–256 (2003).

83 Janušonis, S., Gluncic, V. & Rakic, P. Early serotonergic projections to Cajal-Retzius cells: relevance for cortical development. Journal of Neuroscience 24, 1652–1659 (2004).

84 Dölen, G. Autism: Oxytocin, serotonin, and social reward. Social neuroscience 10, 450–465 (2015).

85 Veenstra-VanderWeele, J. et al. Autism gene variant causes hyperserotonemia, serotonin receptor hypersensitivity, social impairment and repetitive behavior. Proceedings of the National Academy of Sciences 109, 5469–5474 (2012).

86 Olivier, J., Blom, T., Arentsen, T. & Homberg, J. The age-dependent effects of selective serotonin reuptake inhibitors in humans and rodents: A review. Progress in neuro-psychopharmacology and biological psychiatry 35, 1400–1408 (2011).

87 Lesch, K.-P. & Waider, J. Serotonin in the modulation of neural plasticity and networks: implications for neurodevelopmental disorders. Neuron 76, 175–191 (2012).

88 Lesch, K.-P. et al. Association of anxiety-related traits with a polymorphism in the serotonin transporter gene regulatory region. Science 274, 1527–1531 (1996).

89 Fernández, M., Mollinedo-Gajate, I. & Peñagarikano, O. Neural circuits for social cognition: implications for autism. Neuroscience 370, 148–162 (2018).

90 Migliarini, S., Pacini, G., Pelosi, B., Lunardi, G. & Pasqualetti, M. Lack of brain serotonin affects postnatal development and serotonergic neuronal circuitry formation. Molecular psychiatry 18, 1106–1118 (2013).

91 Richardson-Jones, J. W. et al. Serotonin-1A autoreceptors are necessary and sufficient for the normal formation of circuits underlying innate anxiety. Journal of Neuroscience 31, 6008–6018 (2011).

92 Lira, A. et al. Altered depression-related behaviors and functional changes in the dorsal raphe nucleus of serotonin transporter-deficient mice. Biological psychiatry 54, 960–971 (2003).

93 Rood, B. D. et al. Dorsal raphe serotonin neurons in mice: immature hyperexcitability transitions to adult state during first three postnatal weeks suggesting sensitive period for environmental perturbation. Journal of Neuroscience 34, 4809–4821 (2014).

94 Berger, M. A., Barros, V. G., Sarchi, M. I., Tarazi, F. I. & Antonelli, M. C. Longterm effects of prenatal stress on dopamine and glutamate receptors in adult rat brain. Neurochemical research 27, 1525–1533 (2002).

95 Fride, E. & Weinstock, M. Prenatal stress increase anxiety related behavior and alters cerebral lateralization of dopamine activity. Life sciences 42, 1059–1065 (1988).

96 Zill, P. et al. SNP and haplotype analysis of a novel tryptophan hydroxylase isoform (TPH2) gene provide evidence for association with major depression. Molecular psychiatry 9, 1030 (2004).

97 Gao, J. et al. TPH2 gene polymorphisms and major depression-a meta-analysis. PloS one 7, e36721 (2012).

98 Canli, T. & Lesch, K.-P. Long story short: the serotonin transporter in emotion regulation and social cognition. Nature neuroscience 10 (2007).

99 Swedo, S. E. et al. A controlled trial of light therapy for the treatment of pediatric seasonal affective disorder. Journal of the American Academy of Child & Adolescent Psychiatry 36, 816–821 (1997).

100 Niederhofer, H. & von Klitzing, K. Bright light treatment as mono-therapy of non-seasonal depression for 28 adolescents. International journal of psychiatry in clinical practice 16, 233–237 (2012).

101 Krysta, K., Krzystanek, M., Janas-Kozik, M. & Krupka-Matuszczyk, I. Bright light therapy in the treatment of childhood and adolescence depression, antepartum depression, and eating disorders. Journal of Neural Transmission 119, 1167–1172 (2012).

