## Supplemental Figures for "Photoperiodic Effects on Monoamine Signaling & Gene Expression Throughout Development in the Serotonin & Dopamine Systems"

Running Title: **Photoperiod Impacts Monoamine Content & Gene Expression**

Justin K. Siemann[a] PhD, Piper Williams[b], Turnee N. Malik[a], Chad Jackson[a] PhD, Noah H. Green[a] PhD, Ronald Emeson[a] PhD, Pat Levitt[b] PhD, and Douglas G. McMahon[a,1] PhD

[a]Vanderbilt University, Nashville, TN, 37232, [b]Children's Hospital of Los Angeles, Los Angeles, CA, 90027.

[1]Corresponding Author: Douglas G. McMahon, Biological Sciences, Vanderbilt University, 8270 MRB III BioSci Bldg, 465 21st Ave South, Nashville, TN, 37232, USA.

Contact information:, (615) 936-7108

**Supplementary Figures**

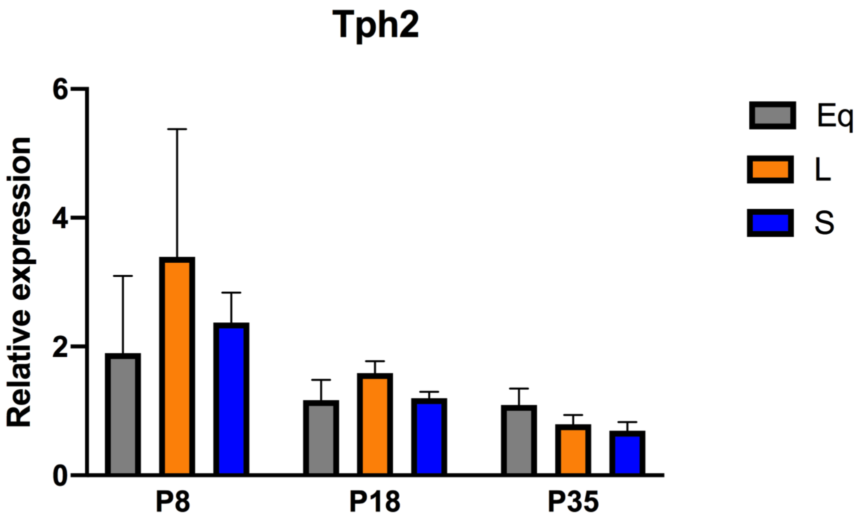


**Supplementary Figure 1.** No significant main effect of photoperiod ($p = 0.6330$; $F (2, 41) = 0.4625$), however a significant main effect of age ($p = 0.0281$; $F (2, 41) = 3.901$) was found for midbrain *Tph2* expression of mice developed under either Equinox (Eq), Long (L), or Short (S) photoperiods. Reference gene for relative expression was β-actin.

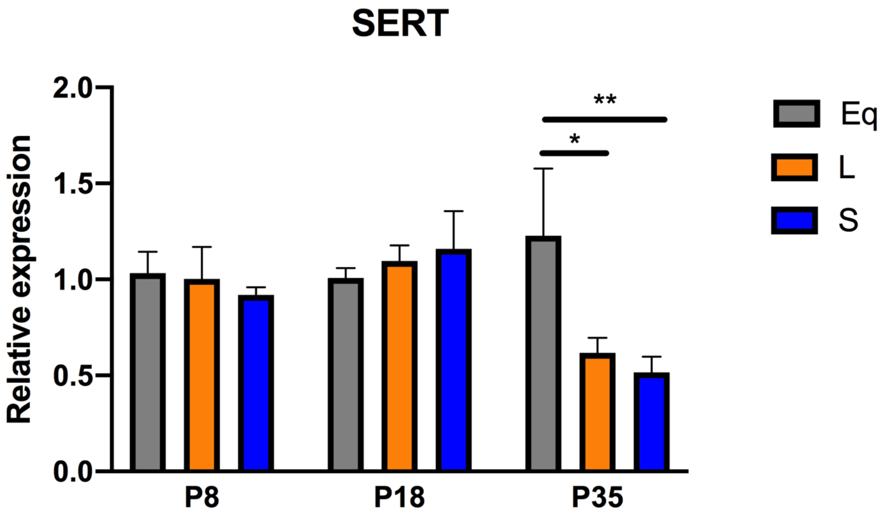


**Supplementary Figure 2.** No significant main effect of photoperiod ($p = 0.1913$; $F (2, 45) = 1.716$) was found for midbrain *SERT* expression for mice developed under either Equinox (Eq), Long (L), or Short (S) photoperiods. Significant post-hoc tests revealed differences at P35 between Equinox and Long ($p = 0.0179$), and Equinox and Short ($p = 0.0078$) photoperiod groups with the significance levels being as follows: (* = $p < 0.05$, ** = $p < 0.01$). Reference gene for relative expression was β-actin.

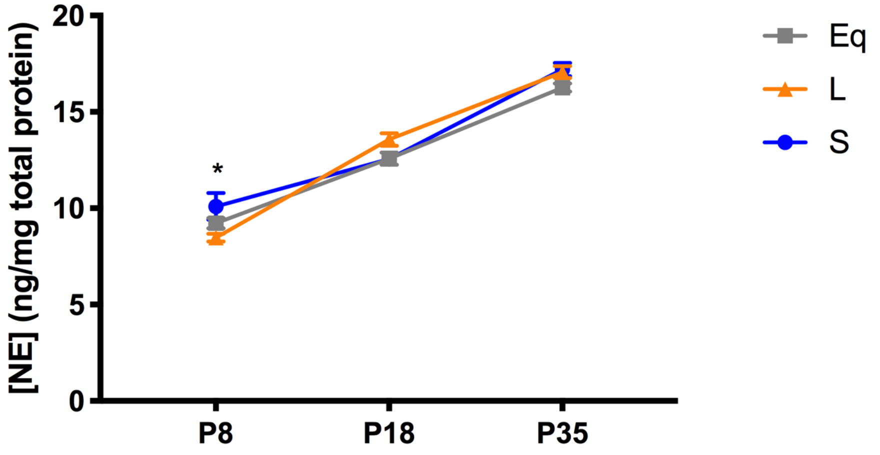


**Supplementary Figure 3.** No significant main effect of photoperiod ($p = 0.1627$; $F(2, 102) = 1.849$), a significant main effect of age ($p < 0.0001$; $F(2, 102) = 287.9$), and a significant interaction effect ($p = 0.0124$; $F(4, 102) = 3.369$) were observed for midbrain norepinephrine content of mice developed under either Equinox (Eq), Long (L), or Short (S) photoperiods. A significant Holm-Sidak's multiple comparison post-hoc test revealed a difference at P8 between Short and Long ($p = 0.0245$) photoperiod groups. The significance level is as follows: (* = $p < 0.05$).

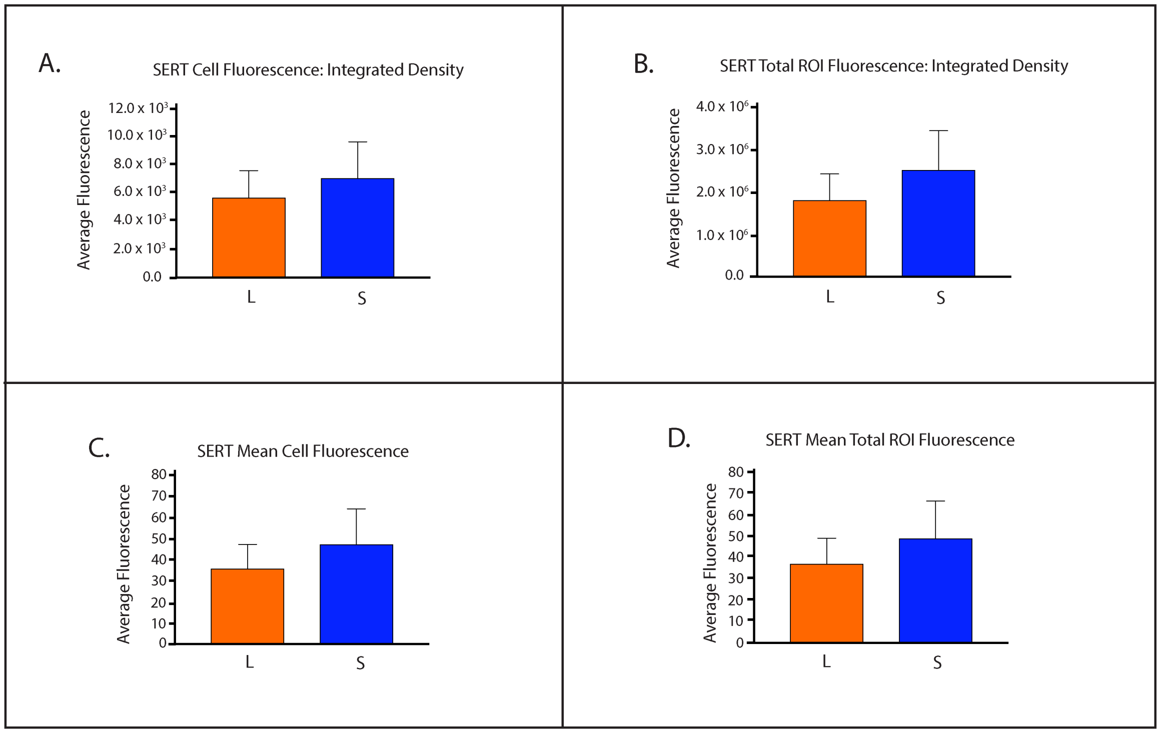


**Supplementary Figure 4.** No significant differences observed in *SERT* expression in the DRN of mice developed under Long or Short photoperiod conditions. **A)** Integrated density of cell fluorescence, **B)** Integrated density of ROI fluorescence, **C)** Quantification of mean cell fluorescence, and **D)** Quantification of ROI mean fluorescence. Mice developed under either Long (L) or Short (S) photoperiods and RNAScope experiments occurred at P50. While no significant differences were observed between the groups, note how *SERT* expression was reduced in all four measures for Long compared to Short photoperiod conditions.

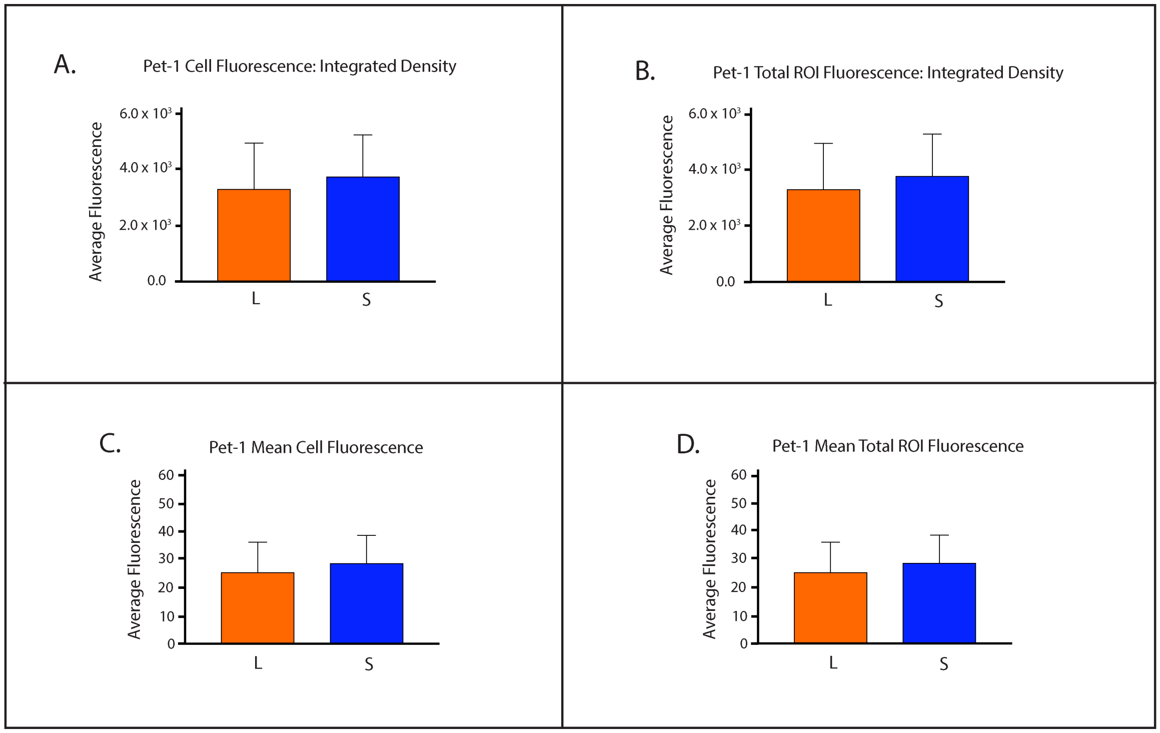


**Supplementary Figure 5.** No significant differences found in *Pet-1* expression in the DRN of mice developed under Short compared to Long photoperiods. **A)** Integrated density of cell fluorescence, **B)** Integrated density of ROI fluorescence, **C)** Quantification of mean cell fluorescence, and **D)** Quantification of ROI mean fluorescence. Mice developed under either Long (L) or Short (S) photoperiods and RNAScope experiments occurred at P50.

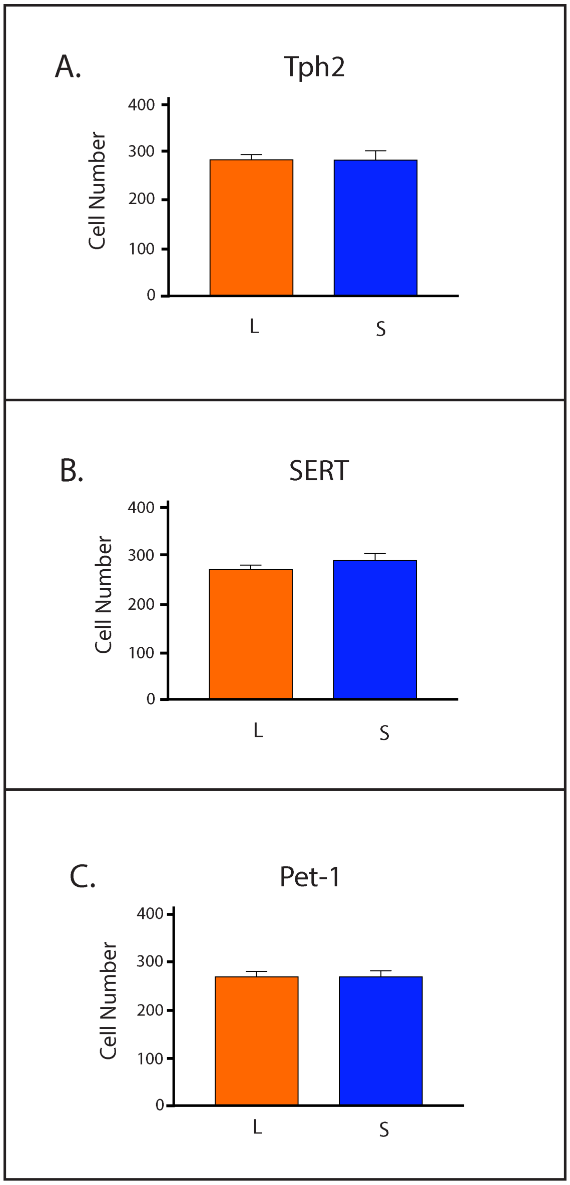


**Supplementary Figure 6.** No significant differences in cell number found for key serotonergic genes in the DRN of C3Hf$^{+/+}$ mice developed under Long or Short photoperiod conditions. Cell number for **A)** *Tph2*, **B)** *SERT*, and **C)** *Pet-1* expression. Mice developed under either Long (L) or Short (S) photoperiods and RNAScope experiments occurred at P50.
